# Endocrine, metabolic, and endocannabinoid alterations in rats exhibiting high anxiety-related behaviors

**DOI:** 10.1101/605329

**Authors:** Julio David Vega-Torres, Priya Kalyan-Masih, Donovan A. Argueta, Nicholas V. DiPatrizio, Johnny D. Figueroa

## Abstract

Anxiety disorders are major risk factors for obesity. However, the mechanisms accounting for this susceptibility remain unclear. Animal models have proved to be useful tools for understanding the role of emotional functioning in the development and maintenance of metabolic alterations implicated in obesity. Here we sought to determine the predictive value of behavioral indices of anxiety for hormonal and metabolic imbalances in rats. Adult Lewis rats were screened on the elevated plus maze (EPM). K-means clustering was used to divide the rats into two groups based on their anxiety index in the EPM: low (LA) and high anxiety (HA) rats. This proxy of anxiety combines individual EPM parameters and accepted ratios into a single score. Four weeks later, we measured markers of endocrine and metabolic function. We found that relative LA rats, the HA rats exhibited reduced latencies to exit a modified light-dark conflict test. Our results show that the HA rats displayed increased corticosterone levels when compared to LA rats. Furthermore, the HA rats weighted more and exhibited an enhanced glycemic response to exogenously administered glucose during the glucose tolerance test, indicating glucose intolerance. Notably, when compared to LA rats, the HA rats showed higher circulating levels of the endogenous cannabinoid, 2-arachidonoyl-sn-glycerol (2-AG). Together, these data indicate that patterns of emotional reactivity associated with anxiety may share common pathological pathways with metabolic complications implicated in obesity. Uncovering metabolic risk factors for anxiety disorders have the potential to strongly impact how we treat mental illnesses.

**Objective:** To test the hypothesis that indices of anxiety in the elevated plus maze have predictive value for alterations implicated in obesity, which include heightened emotional reactivity, and dysregulated corticosterone, glucose metabolism, and circulating endocannabinoid levels.

## Introduction

Anxiety disorders are the most common of mental disorders, affecting more than 30% of adults at some point in their lives (1). Anxiety has been defined as “a psychological, physiological, and behavioral state induced in animals and humans by a threat to well-being or survival, either actual or potential” (2). Anxiety is characterized by the activation of the autonomic and neuroendocrine systems, heightened arousal, and specific behavioral responses, including escape, avoidance, and overall readiness to respond to danger. Although anxiety is a normal reaction to threat or perceived environmental challenges and essential for survival, excessive and maladaptive anxiety can become pathological. Classical theories of causation support that pathological anxiety manifests in vulnerable individuals and can be triggered by psychological factors (3). However, more integrative views of anxiety disorders are also considering the critical contribution of biological factors to the etiology of pathological anxiety.

Anxiety disorders are highly comorbid with metabolic disorders. There is growing recognition for a reciprocal, bidirectional link between anxiety disorders and pro-inflammatory diets (4), and obesity and related metabolic diseases (5; 6). Studies in humans and rodents have shown that obesity and the consumption of obesogenic diets are risk factors for the development of anxiety-related psychopathology (7–10). Interestingly, a new study suggests that these differences in emotional and behavioral responses related to anxiety appear with changes in body weight in obesity-prone rats (11). Other studies demonstrate that individuals exposed to psychosocial stress and diagnosed with anxiety disorders are susceptible to increases in body mass index (BMI) and obesity (12; 13). Although the mechanisms accounting for the link between obesity and anxiety remain speculative, this bidirectionality suggests common pathophysiological processes between physical and mental illnesses. Thus, this experiment was undertaken to examine whether anxiety-like phenotypes are predictive for pathogenic processes implicated in obesity and related metabolic disorders. We hypothesized that rats exhibiting higher anxiety-like behaviors will share endocrine and metabolic traits comparable to obesity.

## Results and Discussion

There is a growing appreciation for the substantial individual differences in anxiety-like behaviors in rats. The elevated plus maze (EPM) have proved to be a successful method to systematically determine individuality in anxiety-like and avoidance behaviors (14–16). However, to our knowledge, no studies have investigated whether individual differences in anxiety-like behaviors in the EPM may predict endocrine and metabolic changes co-occurring with obesity in non-obese rats. We began our investigation of the association between anxiety and obesity-like phenotypes by hypothesizing that proxies of anxiety in the EPM may have predictive value for markers of dysregulated metabolism. We used the k-means clustering algorithm to group rats in two groups based on their anxiety-like behaviors in the EPM. As expected, rats in the low anxiety (LA) group exhibited reduced indices of anxiety in the EPM relative to the high anxiety (HA) group. Our analyses revealed that rats from the HA group had a significant 29% increase in anxiety index (AI) when compared to LA group (HA: 0.63 ± 0.02 vs. LA: 0.49 ± 0.01; *t*_(16)_ = 4.82, *p* = 0.0002) (Figure 1B). This finding is confirmed in the track plot shown in Figure 1C, demonstrating that HA rats exhibited high avoidance of the open arms while spending most of the time in the closed arms of the EPM. Although HA rats exhibited higher acoustic startle responses, we found no significant differences between groups (data not shown).

**Figure 1.**
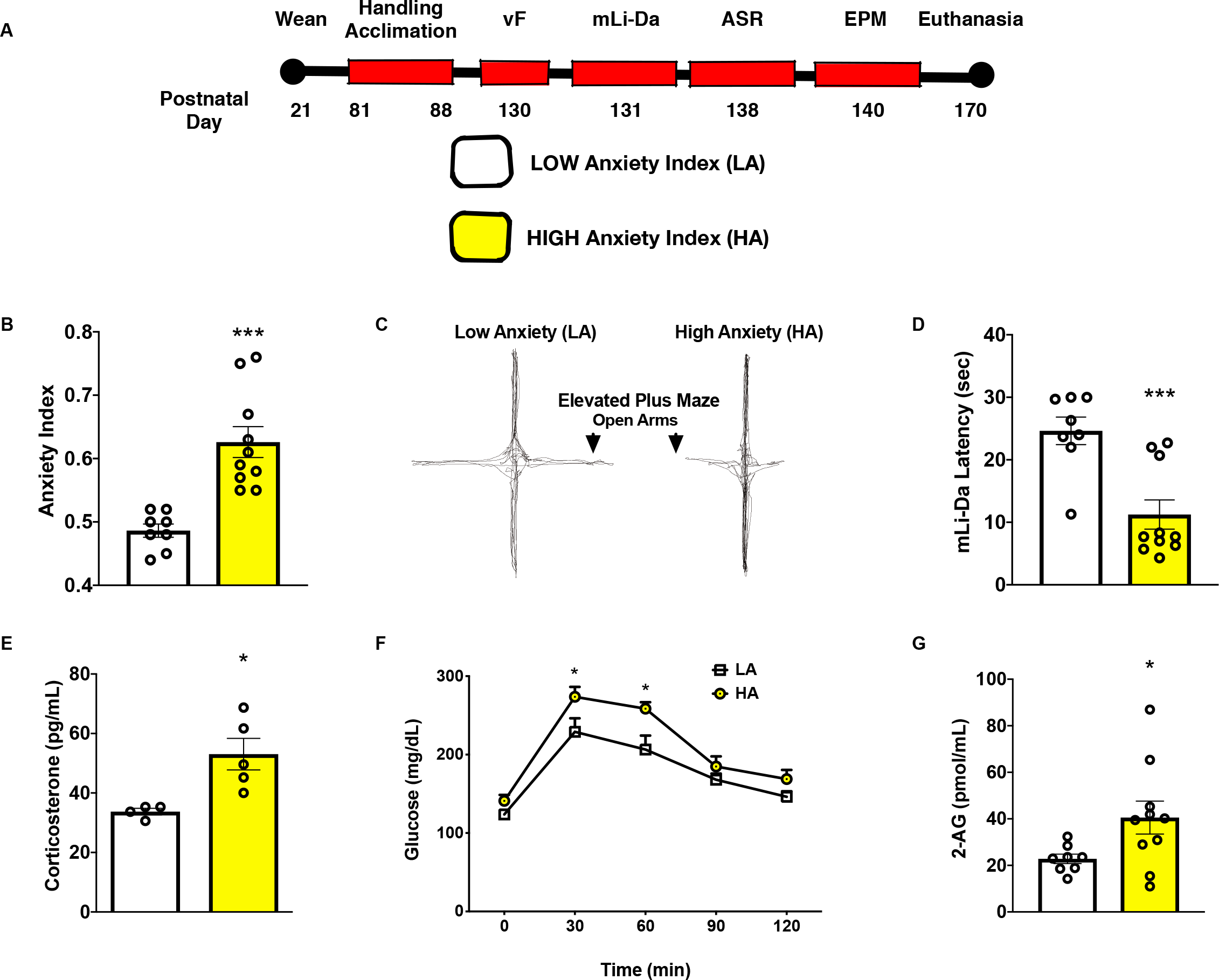
High-anxiety rats exhibit increased open arm avoidance while manifesting disruptions in levels of corticosterone, glucose, and 2-AG. **(A)** Timeline of the study showing the order of the various behavioral tests performed. **(B)** Anxiety index of LA and HA rats following k-means clustering. HA rats exhibited higher indices of anxiety in the EPM (29% increase) when compared to LA rats (\*\*\**p* < 0.001). **(C)** Representative track plot of the elevated plus maze (EPM) for LA and HA rats. These plots highlight that HA rats were avoiding the open arms while spending most of the time in the closed arms of the EPM. **(D)** Average latency obtained from the modified light-dark box behavioral test for LA and HA rats. HA rats had a significant reduction (54%) in latency (time to exit the aversive light chamber) relative to LA rats (\*\*\**p* < 0.001). **(E)** Average levels of corticosterone in feces. HA rats exhibited a significant increase (57%) in fecal corticosterone levels when compared to LA (\**p* < 0.05). **(F)** Average glucose concentration at baseline, and 30 min, 60 min, 90 min, and 120 min following intraperitoneal glucose tolerance test. HA rats exhibited increased glucose levels at 30 min (20% increase; \**p* > 0.05) and at 60 min (26% increase; \**p* < 0.05) when compared to LA rats. **(G)** Average levels of 2-AG in plasma. HA rats had a significant increase (61%) in circulating levels of 2-AG relative to LA rats (\**p* < 0.05).

The conflict-avoidance test was used to evaluate others measures of anxiety-like behaviors associated with the rats’ innate light aversion (photophobia). In this test, the rats are first trained to escape the aversive bright light chamber and navigate through a 3 mm sharp probe chamber to get to a less aversive dark chamber. This test has been used to evaluate nociception, and cognitive and motivational processing (17). Our analyses showed that HA rats had a significant reduction (54%) in latency (time to exit light chamber after door is opened) when compared to LA rats (LA: 24.63 ± 2.2 vs. HA: 11.24 ± 2.34; *t*_(16)_ = 4.09, *p* = 0.0009) (Figure 1D), indicating heightened avoidance behaviors and validating the EPM-based group classification. We used an electronic von Frey anesthesiometer to determine nociceptive thresholds in the LA and HA rats. We reasoned that this test would provide information as to whether differences in escape latencies were related to reduced nociception in the HA rats. Two-way ANOVA analyses revealed no significant group [*F*_(1, 16)_ = 0.01, *p* = 0.92] effects on withdrawal threshold in neither limb (data not shown), indicating similar nociceptive thresholds between groups.

Corticosterone is a common marker of anxiety- and stress-related behaviors in rats and implicated in the detrimental effects of obesogenic diets on emotionality (9; 18; 19). We measured fecal corticosterone levels at the end of the study (30 days after EPM testing). Interestingly, the HA rats exhibited a significant increase (57%) in fecal corticosterone levels when compared to LA (LA: 33.73 ± 1.1 vs. HA: 53.05 ± 5.3; *t*_(7)_ = 3.17, *p* = 0.01) (Figure 1E). Supporting the well-established link between corticosterone, anxiety, and glucose metabolism, we found that HA rats showed impairments in glucose metabolism. Analyses revealed significant group [*F*_(1, 9)_ = 9.88, *p* = 0.012] and time [*F*_(4, 36)_ = 46.94, *p* < 0.0001] effects, while no significant interactions [*F*_(4, 36)_ = 1.27, *p* = 0.30] effects on glucose clearance (Figure 1F). Sidak’s post-hoc analyses showed that, relative to LA rats, the HA rats exhibited increased glucose levels at 30 min (20% increase; *p* = 0.044) and at 60 min (26% increase; *p* = 0.013) following the bolus glucose solution injection. Subsequently, we sought to determine whether proxies of anxiety correlate with changes in body weight. We found increased body mass in HA rats relative to LA rats. Two-way ANOVA analyses revealed significant group [*F* <_(1, 16)_ = 7.14, *p* = 0.017] and time [*F*_(2, 32)_ = 2,721, *p* < 0.0001], while no significant interactions [*F*_(2, 32)_ = 2.27, *p* < 0.119] effects on body weight (data not shown). Notably, these differences in body weight emerged at the last time point evaluated (6% increase at PND 170; *p* = 0.006) and before or during the week that the behaviors were recorded. A feature common to changes in obesity is alterations in leptin, a major orchestrator of metabolism and regulator of emotionality (20). Our analyses revealed that rats from HA group had a significant increase in plasma leptin levels when compared to LA group (HA: 1364 ± 237.4 vs. LA: 638.3 ± 121.4; *t*_(11)_ = 2.58, *p* = 0.025) (data not shown). Together, these data support the relationship between body weight, circulating leptin levels, and glucose homeostasis and its potential association with anxiety-related phenotypes.

In search of a unifying biological factor for the above results, we investigated the levels of major circulating endocannabinoids. Several lines of evidence show that the endocannabinoid system influences a wide variety of neurobiological processes including emotionality, cognitive function, and glucocorticoid-mediated stress responses. This system is an attractive area for research into new pharmacotherapeutic targets for anxiety disorders (21). Studies show that both reductions (22) or elevations (23) in circulating endocannabinoid levels in patients diagnosed with post-traumatic stress disorder (PTSD), suggesting that dysregulated endocannabinoid signaling contributes to mental illnesses. The endocannabinoid system in the periphery has gained recent attention for its roles in controlling feeding behavior and energy metabolism (24; 25). For example, endocannabinoid levels increase in the upper small-intestinal epithelium of rodents after tasting dietary lipids (26; 27) fasting (28–30), and are elevated in blood in rodent and human diet-induced obesity (29; 31–34). Furthermore, inhibiting peripheral cannabinoid receptor 1 (CB_1_R) blocks sham feeding of dietary fats (26; 27), refeeding after a 24-h fast (30), and hyperphagia associated with diet-induced obesity (33). In humans, hedonic eating was also associated with higher levels of 2-AG in circulation (35; 36). In the present study, we find increases in circulating levels of the endocannabinoid, 2-AG, in rats exhibiting a “high anxiety” phenotype (61% increase relative to LA; LA: 22.81 ± 2.03 vs. HA: 40.54 ± 7.09; *t*_(16)_ = 2.17, *p* = 0.046) (Figure 1G). This result raises the intriguing possibility that increased levels of 2-AG under these conditions may impact feeding and other behaviors. Furthermore, diet-induced changes in 2-AG levels may influence affect and emotionality, resulting in anxiety-related phenotypes. However, a direct test of these hypotheses remains for future studies.

## Conclusions

We found significant alterations in obesity-related endocrine and metabolic biomarkers based on individual differences in anxiety-like behaviors. Collectively, our findings confirm that anxiety-like behaviors share common pathogenic pathways with endocrine and metabolic complications of obesity. This model may present new opportunities in the prevention and amelioration of co-morbid obesity and psychiatric disorders.

### Limitations

Our results indicate that HA Lewis rats exhibit metabolic, hormonal, and endocannabinoid alterations consistent with obesogenic phenotypes. However, it is difficult to dissociate whether the observed changes enhance susceptibility to an anxiety-like phenotype, or whether the anxiety-like behavioral manifestations in the EPM result in metabolic alterations. Replication in larger cohorts, and with different rat strains and diets is warranted. Considerations for sex as a biological variable should be factored in future studies.

### Conjectures

While we observed distinctive behavioral, endocannabinoid, metabolic, and hormonal profiles in rats exhibiting high anxiety-like behaviors in the elevated plus maze, it is unclear from our data whether the neuroendocrine and metabolic changes accompanying weight gain promote anxiety or vice versa. Future studies are required to test a causal relationship. Importantly, further research is also required to estimate whether the metabolic-related alterations in rats exhibiting high indices of anxiety can be targeted to ameliorate both, anxiety- and obesity-like phenotypes.

## Methods

Experimental procedures were performed in compliance with regulations and institutional guidelines consistent with the National Institutes of Health *Guide for the Care and Use of Laboratory Animals*. Fifty-four (54) adolescent male Lewis rats were purchased from Charles River Laboratories (Portage, MI). The rats were grouped (2 per cage) and fed *ad libitum*. Rats were fed purified diets based on standard AIN-93G formulations (7 g% fat; mainly from soybean oil). The rats were kept under standard housing conditions (22 ± 2 °C and regular 12 h dark/light cycle). Animals were handled for 5 days before behavioral testing. Behavioral tests were conducted between 10:00-14:00 h.

### Elevated Plus Maze (EPM)

The rats were tested on the EPM at PND 130. The near infra-red-backlit EPM consisted of two opposite open arms (50.8 × 10.2 cm) and two enclosed arms (50.8 × 10.2 × 40.6 cm) elevated 72.4 cm above the floor (MedAssociates, St. Albans, VT, USA) The junction area between the four arms measured 10 × 10 cm. Behaviors were recorded in a dark room. The rats were placed on the central platform facing an open arm at the beginning of each 5-min trial. The maze was cleaned after each test session. Arm activity (total number of entries and duration in each arm) was determined using Ethovision^®^ XT tracking software. From this data, we calculated the well-validated anxiety index according to Cohen et al., (37; 38) and Contreras et al., (39):

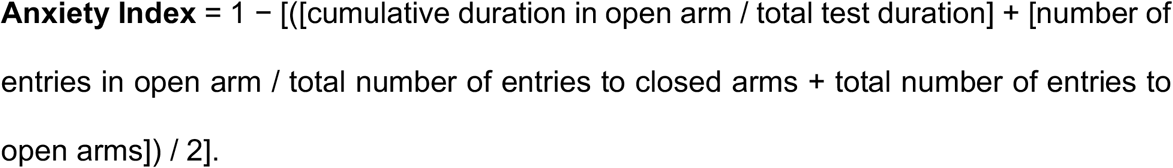

### Acoustic Startle

The acoustic startle reflex (ASR) were performed using the SR-Lab Acoustic Chambers (San Diego Instruments). An experimental session began with a habituation program, consisting of 5 min of background noise at 68 dB. Following habituation, the initial block consisted of 11 acoustic startle trials was recorded (120 dB; 20 ms). The trials were presented in a randomized manner (intertrial interval = 10 –25 s). The second block had 36 trials presented in a pseudorandomized order. In this block, the trials included pulse-alone trials (120 dB) and pre-pulse trials (five trials each: 71, 74, or 80 dB; 40 ms duration). The third block of the session consisted of 5 additional pulse-alone startle trials (120 dB). Subsequently, the rats were returned to their cages and each chamber was cleaned with soap and water and thoroughly dried. Max startle amplitudes during the session were averaged.

### Modified Light-Dark Test: Mechanical Conflict-Avoidance System

Conflict-avoidance behaviors were measured using the Mechanical Conflict-Avoidance System (MCS) (Noldus). The MCS consisted of one light and one dark chamber (16.5 cm wide × 21.5 cm deep × 15.25 cm high, each) connected by an enclosed alleyway chamber (39.5 cm wide × 21.5 cm deep × 15.25 cm high). All three chambers were constructed from acrylic resin and colored red. An array of stainless-steel sharp nociceptive probes (tip dimension = 0.4 mm) was embedded below the floor of the alleyway chamber. The probe array was lowered below the alleyway floor for training sessions and elevated for testing. In this system, anxiety-like behaviors are expressed as decreased escape latency from bright light to the dark compartment. The MCS protocol consisted of 5 days. We used a dim red-light lamp for the whole duration of the experiments. During the first two days, the rats were acclimated to the MCS. The light of the device was turned off and the rats placed in the light chamber (LC). After 10 seconds, the aversive bright light was turned on and remained on for 20 seconds. Afterwards, the LC guillotine door was carefully opened and the rats allowed to explore the MCS for 4.5 minutes (total duration of each acclimation session was ~5 minutes). The experimental sessions took place on days 3 to 5. During each testing session, different probe heights were used on each day (0 mm for *day 3*, 1 mm for *day 4*, and 3 mm for *day 5*). For testing days, the light of the device was turned off and the rat was placed in the LC. After 10 seconds, the light was turned on and remained on for 20 seconds. The LC door was carefully opened and the time to exit the LC was recorded. Each rat was given 30 seconds to exit LC which was counted as the latency time. After the rats spent 5-10 seconds in the dark chamber, the lid was removed and the rats were returned to their home cages. The apparatus was clean thoroughly between each test. For each experimental session, we repeated this process 3 times per rat with at least 10 minutes between trials.

### Von Frey

Sensory thresholds to a mechanical stimulus were measured using an electronic von Frey aesthesiometer (IITC Life Science, Woodland Hills, CA). Each rat was placed in a Plexiglas chamber on top of an elevated mesh stand that provided access to the plantar surface of the paws. The rats were allowed to acclimate to the testing chamber for 30 min prior to testing. Three mock trials were performed halfway through the acclimation period. Subsequently, a rigid blunt tip that was attached to the aesthesiometer was applied to the plantar surface. The withdrawal threshold or mechanical sensitivity was defined as the average force (g) required for paw withdrawal in five trials separated by at least 2 min interval. The maximum and minimum threshold values were excluded from each paw before analyses.

### Intraperitoneal glucose tolerance test (IPGTT)

The IPGTT was performed at PND 170. The rats were fasted for 12 h prior to being anesthetized with pentobarbital (100 mg/kg). Subsequently, the rats received intraperitoneal injections of 40% dextrose solution (2 g of dextrose per kg of body weight;). Blood was sampled at 0, 30, 60, 90, and 120 min for glucose analyses. Glucose levels were measured using a glucometer (OneTouch UltraMini^®^ manufactured by LifeScan, Inc Milpitas, CA, USA).

### Measuring corticosterone and leptin concentrations using ELISA

The rats were placed on new bedding 24 h before feces collection. Fecal samples were collected from each cage at PND 170 and corticosterone extracted following previous methods (40). Briefly, fecal corticosterone was extracted by incubating feces in ethanol overnight at a ratio of 10 mL ethanol per gram of feces. Corticosterone concentration was determined using the Correlate EIA kit (Assay Designs, Ann Arbor, MI). ELISA plates were read at 405 nm (Qcal Biotek Instruments plate reader, Winooski, VT USA). Specific concentrations for each sample were determined as percentage bound using a standard curve of samples ranging from 32 to 20,000 pg/mL. Values were then calculated and reported as picograms of corticosterone per mL.

Leptin plasma levels were measured using a commercial ELISA kit (Abcam). Plasma samples were diluted with kit assay buffer (1:3 dilutions). ELISA plates were read at 450 nm on a plate reader (Molecular Devices). Specific concentrations for each sample were determined as mean absorbance using a standard curve of samples ranging from 0 to 8000 pg/ml.

### Lipid extraction

2-AG was extracted from plasma according to published methods (33; 41). Blood was collected by cardiac puncture and stored in tubes with EDTA, then plasma was obtained by centrifugation (1500 *g* for 10 minutes, maintained at 4°C). All samples were stored at −80°C until processing, at which time plasma (0.1 mL) was added to 1.0 mL of methanol solution containing the internal standard, [^2^H_5_] 2-AG (Cayman Chemical, Ann Arbor, MI, USA). Lipids were extracted with chloroform (2 mL) and washed with 0.9 % saline (0.9 mL). Organic phases were collected and separated by open-bed silica gel column chromatography as described (42), then dried under N_2_ stream (99.998% pure) and resuspended in 0.1 mL of methanol:chloroform (9:1). Quantitation of 2-AG levels was made by ultra-performance liquid chromatography coupled to tandem mass spectrometry (UPLC/MS/MS), detailed below.

### Measurement of 2-AG

Data was collected using an Acquity I Class UPLC system coupled to a Xevo TQ-S Mass Spectrometer (Waters, Milford, MA, USA) with accompanying electrospray ionization (ESI) interface. Lipids were separated on an Acquity UPLC BEH C_18_ column (2.1 × 50 mm i.d., 1.7µm, Waters) with inline Acquity guard column (UPLC BEH C_18_ VanGuard Pre-column; 2.1 × 5 mm i.d., 1.7µm, Waters), and eluted by a gradient of methanol in water (0.25% acetic acid, 5mM ammonium acetate) according to the following gradient at a flow rate of 0.4 mL per min: 80% methanol 0.5 min, 80% to 100% methanol 0.5-2.5 min, 100% methanol 2.5-3 min, 100% - 80% methanol 3-3.1 min). Column temperature was maintained at 40° C, and samples were maintained in the sample manager at 10° C. Argon (99.998%) was used as collision gas. MS detection was in positive ion mode and capillary voltage set at 0.1 kV. Cone voltage and collision energy as follows, respectively: 2-AG = 30v, 12v; [^2^H_5_] 2-AG = 25v, 44v. Lipids were quantified using a stable isotope dilution method detecting protonated adducts of the molecular ions [M+H]^+^ in the multiple reaction monitoring (MRM) mode. Extracted ion chromatograms were used to quantify 2-AG (*m/z* = 379.3>287.3) and [^2^H_5_] 2-AG (*m/z* = 384.3 > 93.4), which was used as an internal standard.

### Statistics

Statistical analysis was performed using GraphPad Prism 8. We used the K-means clustering method to assign all rats to two groups according to their anxiety index (SPSS Version 25.0; IBM Corp., Armonk, NY). Statistical differences were evaluated using 2-tailed unpaired Student’s t test or 2-way ANOVA with multiple comparisons, as indicated. P values of less than 0.05 were considered statistically significant.

## Funding Statement

This work was partly supported by Seed Grant Funds to Johnny D. Figueroa, and National Institutes of Health grants MD006988 and GM060507 to Marino De Leon and DA034009, DK119498, and DK114978 to Nicholas V. DiPatrizio.

## Conflict of Interest

The authors declare no conflict of interest.

## Ethics Statement

No fraudulence was committed in performing these experiments or during data analyses. We understand that any case of fraudulence can result in the study being retracted.

